# GPCR signaling promotes severe stress-induced organismic death in *C. elegans*

**DOI:** 10.1101/2022.09.16.508223

**Authors:** Changnan Wang, Yong Long, Bingying Wang, Chao Zhang, Dengke K. Ma

## Abstract

How an organism dies is a fundamental yet poorly understood question in biology. An organism can die of many causes, including stress-induced phenoptosis, also defined as organismic death that is regulated by its genome-encoded programs. The mechanism of stress-induced phenoptosis is still largely unknown. Here we show that transient but severe freezing-thaw stress (FTS) in *C. elegans* induces rapid and robust phenoptosis that is regulated by G-protein coupled receptor (GPCR) signaling. RNAi screens identify the GPCR-encoding *fshr-1* in mediating transcriptional responses to FTS. FSHR-1 increases ligand interaction upon FTS and activates a cyclic AMP-PKA cascade leading to a genetic program to promote organismic death under severe stress. FSHR-1/GPCR signaling up-regulates the bZIP-type transcription factor ZIP-10, linking FTS to expression of genes involved in lipid remodeling, proteostasis and aging. A mathematical model suggests that genes may promote organismic death under severe stress conditions, potentially benefiting growth of the clonal population with individuals less stressed and more reproductively privileged. Our studies reveal roles of FSHR-1/GPCR-mediated signaling in stress-induced gene expression and phenoptosis in *C. elegans*, providing empirical new insights into mechanisms of stress-induced phenoptosis with evolutionary implications.

## INTRODUCTION

Organisms of all life forms on earth live with varying but limited lifespans. While the natural aging process and regulation of lifespan in many different organisms have been extensively studied (Antebi 2007; Fontana & Partridge 2015; Kenyon 2010; López-Otín et al. 2013; Mutlu et al. 2021; Shore & Ruvkun 2013), little is known about how a multicellular organism dies at the end of its life after normal aging or in the middle of its life. In nature, the death of an organism can have various causes, including severe abiotic stresses (e.g. heat or starvation), pathogens or predators from ever-changing environments. Prolonged severe abiotic stresses can lead to the death of organisms, in many cases likely because of failure in adapting to the stress, or alternatively through stress-induced phenoptosis (Skulachev 1999; Skulachev 2002; Longo et al. 2005; Skulachev 2019). Using the nematode *C. elegans* as a model organism, we have previously discovered that severe stress from cold shock (CS) followed by warming promotes phenoptosis through a gene transcriptional pathway (Jiang et al. 2018). Discovery of such rapid thermal stress-induced phenoptosis and the genetic tractability of *C. elegans* afford unprecedented access and opportunities to identify the underlying cell signal transduction pathway and mechanisms of stress-induced phenoptosis.

With its small body size, *C. elegans* experiences varying temperature depending on the ambient environment and exhibits stereotypic behavioral and cell physiological response to chronic or acute mild (10-15 °C) hypothermia (Al-Fageeh & Smales 2006; Garrity et al. 2010; Xiao et al. 2013). *C. elegans* can live and reproduce normally within 15-25 °C and increases lifespan when grown under chronic mild hypothermia (15 °C) (Xiao et al. 2013; Lee et al. 2019). Temperature beyond this range causes stress, sterility and organismic death. By studying *C. elegans* mutants abnormally responding to severe thermal (4 °C CS, recovery at 20 °C) stress stimuli, we identified ZIP-10, a bZIP-type transcription factor that promotes severe thermal stress-induced genes and phenoptosis (Jiang et al. 2018). We showed that ZIP-10 activates a genetic program and up-regulates protease-encoding genes in intestinal cells to promote phenoptosis (Jiang et al. 2018). Both the cold-induced *zip-10* expression and the death-promoting effect of ZIP-10 appeared more prominent in old adults than larvae (Jiang et al. 2018). We postulate that genetic programs underlying stress-induced phenoptosis may have evolved because of antagonistic pleiotropic effects and/or “kin selection” at the population level so that the phenoptosis of adult and weak individuals after severe cold-thermal stress would benefit less stressed and reproductively more privileged ones to facilitate the spreading of genes by fit individuals under resource-limiting and high-stress conditions (Jiang et al. 2018; Hamilton 1963; Smith 1964). While evolutionary mechanisms and significance of phenoptosis still remain debatable in the field (Galimov et al. 2019; Kirkwood & Melov 2011; Longo et al. 2005; Sapolsky 2004; Skulachev 2019), many empirical studies suggest that organisms with clonal population structures such as *C. elegans* may indeed chronically age or die under stresses by mechanisms (terminal investment, antagonistic pleiotropy, consumer sacrifice or disposable soma) that can benefit its progeny or reproductive success of the organism itself (Ezcurra et al. 2018; Galimov & Gems 2020; Gulyas & Powell 2021; Wu et al. 2022).

Despite critical roles of ZIP-10 in cold-thermal stress-induced phenoptosis in *C. elegans*, its upstream regulators linking severe stress to the ZIP-10-dependent genetic program remain unidentified. By transcriptome profiling, RNAi screens and genotype-to-phenotype mechanistic analysis, we uncovered a GPCR (FSHR-1, follicular stimulating hormone receptor related) cascade that activates ZIP-10-dependent transcriptional response to freeze-thaw stress (FTS). Activation of this pathway promotes organismic phenoptosis under severe FTS conditions, more prominently in adults than young larval animals. Our findings reveal an essential role of FSHR-1 and its GPCR signaling cascade in stress-induced gene expression, providing mechanistic understanding of phenoptosis and empirical evidence with implications for the evolutionary kin selection theory and the “disposable soma” hypothesis of organismic aging.

## RESULTS

### RNAseq identifies transcriptomic changes to freeze-thaw stress in *C. elegans*

By exploring various severe hypothermic conditions in *C. elegans*, we identified an environmental stress scheme (−20 °C freezing for 45 mins followed by recovery at 20 °C for 24 hrs, FTS) that triggers robust ZIP-10-dependent phenoptosis with higher penetrance than a previous scheme using 4 °C cold shock (Jiang et al. 2018) (Figure 1a). We measured the survival rates of both wild-type and *zip-10* mutants to quantify organismic phenoptosis under such scheme for *C. elegans* at different developmental stages (larval L1, L4, young and older adults). We found that wild-type animals exhibited progressively increased rates of FTS-induced death along with age, with the young (48 hrs post L4) and older adult-stage (5 days post L4) populations reaching nearly 100% death rates after FTS treatment while many larvae survived (Figure 1b). By contrast, FTS-induced phenoptosis was markedly attenuated in *zip-10* mutants, particularly at young adult stages. Given the role of ZIP-10 in organismic death and its regulation by severe cold-thermal stress conditions (Jiang et al. 2018) (and see below), we refer to such severe cold/freezing thermal stress-induced organismic deaths as phenoptosis for simplicity (yet mechanistically do not distinguish between active killing and increased sensitivity). These results establish a robust FTS scheme to induce rapid and environmental stress-triggered phenoptosis in *C. elegans* and also confirm a critical role of ZIP-10 in severe cold/freezing thermal stress-induced phenoptosis.

**Figure 1:**
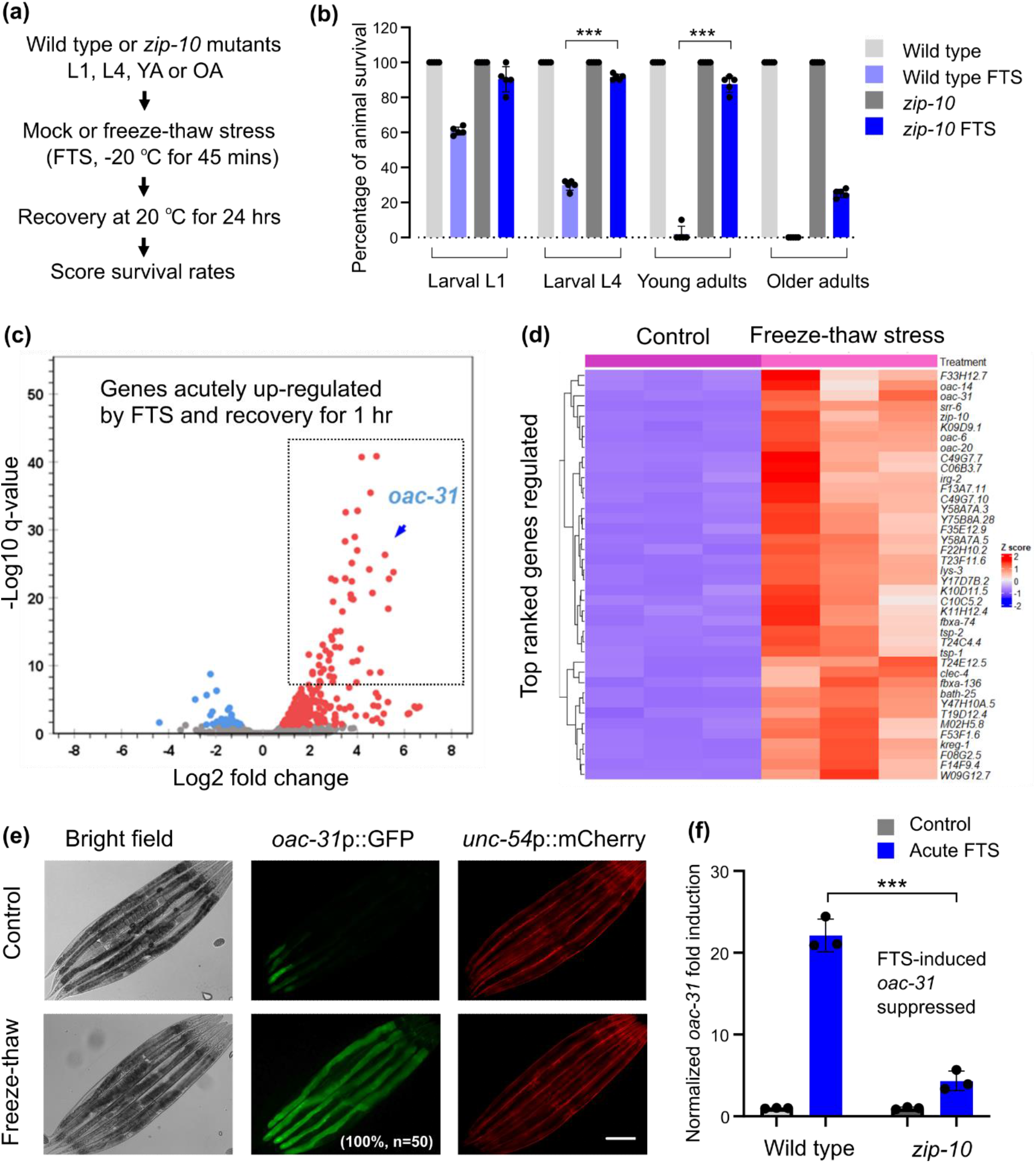
RNAseq identifies transcriptomic changes to freeze-thaw in *C. elegans*. **(a)**, Schematic of experimental flow to assay FTS-induced phenoptosis. **(b)**, Quantification of phenoptosis based on percentages of animals survived after FTS in each condition (young adults, 2 days post L4; older adults, 5 days post L4). Values are means ± S.D with \*\*\**P* < 0.001 (one-way ANOVA with post-hoc Tukey HSD, N = 5 independent experiments, n > 50 for each experiment). **(c)**, Volcano plot showing identified genes that are regulated by FTS based on RNAseq. Red: up-regulated. Blue: down-regulated. **(d)**, Heat map and hierarchical clustering analysis showing the top 50-ranked genes regulated by FTS. **(e)**, Representative epifluorescence images showing *oac-31*p::GFP but not co-injection marker *unc-54*p::mCherry induction by FTS. **(f)**, qRT-PCR showing induction of endogenous *oac-31* by FTS in wild type but attenuated in *zip-10*(*ok3462*) loss-of-function mutants. Values are means ± S.D with \*\*\**P* < 0.001 (two-way ANOVA for genotype-effect interaction and post-hoc Tukey HSD, N = 3 independent experiments, n > 50 for each experiment). Scale bar, 100 μm.

To identify genes acutely regulated by FTS in *C. elegans* that may participate in phenoptosis, we conducted RNA sequencing (RNAseq) of wild-type *C. elegans* populations after 25-minute exposure to −20 °C freezing followed by recovery at 20°C for 1 hr. We used this regime to identify genes that specifically and rapidly respond to FTS rather than those that respond to general organismic deterioration once phenoptosis occurs. After differential expression analyses of triplicate samples, we identified 277 genes that are significantly up- or down-regulated by FTS conditions (Figure 1c-d). Tissue enrichment analysis (TEA, Wormbase) revealed that intestine is the most prominent site of gene regulation by FTS (Figure S1a). FTS-regulated genes do not apparently overlap with those regulated by common stress-responding transcription factors DAF-16 (nutritional stress), SKN-1 (oxidative stress), HSF-1 (heat stress) or HIF-1 (hypoxic stress), and are involved in biological processes including lipid remodeling (e.g. *oac-31, oac-14* and *oac-6*), responses to abiotic and pathogen stresses (e.g. *kreg-1, irg-1* and *lys-3*) (Figure S1b-d). We generated transgenic *C. elegans* strains in which GFP is driven by promoters of the top-ranked FTS-inducible genes. *oac-31*p::GFP emerged as a robust FTS-inducible reporter with low baseline expression and high-fold high-penetrance induction by FTS (Figure 1e). We confirmed by quantitative RT-PCR (qRT-PCR) that FTS up-regulates endogenous *oac-31* in a ZIP-10 dependent manner (Figure 1f). *oac-31* encodes a *C. elegans* homolog of sterol O-acyltransferases and is also the most up-regulated member among the *oac* gene family (Figure S1g). These results identify transcriptomic changes to FTS and led to *oac-31*p::GFP as a robust transcriptional reporter for the genetic response to FTS.

### RNAi screens identify FSHR-1/GPCR signaling in regulating responses to FTS

With an integrated *oac-31*p::GFP transgenic reporter strain, we performed RNAi (Figure 2a) and forward genetic screens to identify genes that are required for FTS induction of *oac-31*p::GFP. For the RNAi screen, we assembled a library of RNAi clones that target genes with adequate intestinal expression (Transcript per million reads, TPM > 2.0) and encoding transmembrane receptors or transcription factors representing a comprehensive set of signal transduction pathways in *C. elegans* (Table S1; Figure S2a). We found that two gene RNAi clones (*sbp-1* and *tra-1*) activated baseline *oac-31*p::GFP, whereas the other two gene RNAi clones (*fshr-1* and *zip-10*) decreased FTS-induced *oac-31*p::GFP (Figure S2b-c). *fshr-1* encodes a glycoprotein hormone receptor homolog in *C. elegans* and has been implicated in mediating oxidative and innate immune responses (Cho et al. 2007, p.1; Kim & Sieburth 2020, p.; Powell et al. 2009; Miller et al. 2015; Kudo et al. 2000). A genetic deletion allele *ok778* (back-crossed to N2 for 5 times) of *fshr-1* fully recapitulated the RNAi phenotype in blocking FTS-induced *oac-31*p::GFP (Figure 2b). Neither FTS nor *fshr-1* RNAi affected *unc-54*p::mCherry, the transgenic co-injection marker for *oac-31*p::GFP. By qRT-PCR, we confirmed that FTS induction of *oac-31* (and also another gene *ncr-1* identified from RNAseq) requires *fshr-1* (Figure S2d). However, at least two other genes, *W09G12.7* and *tsp-1*, do not apparently require *fshr-1* for induction by FTS (Figure S2e). These results reveal an essential and specific role of *fshr-1* in transcriptional up-regulation of FTS-induced genes including *oac-31*.

**Figure 2:**
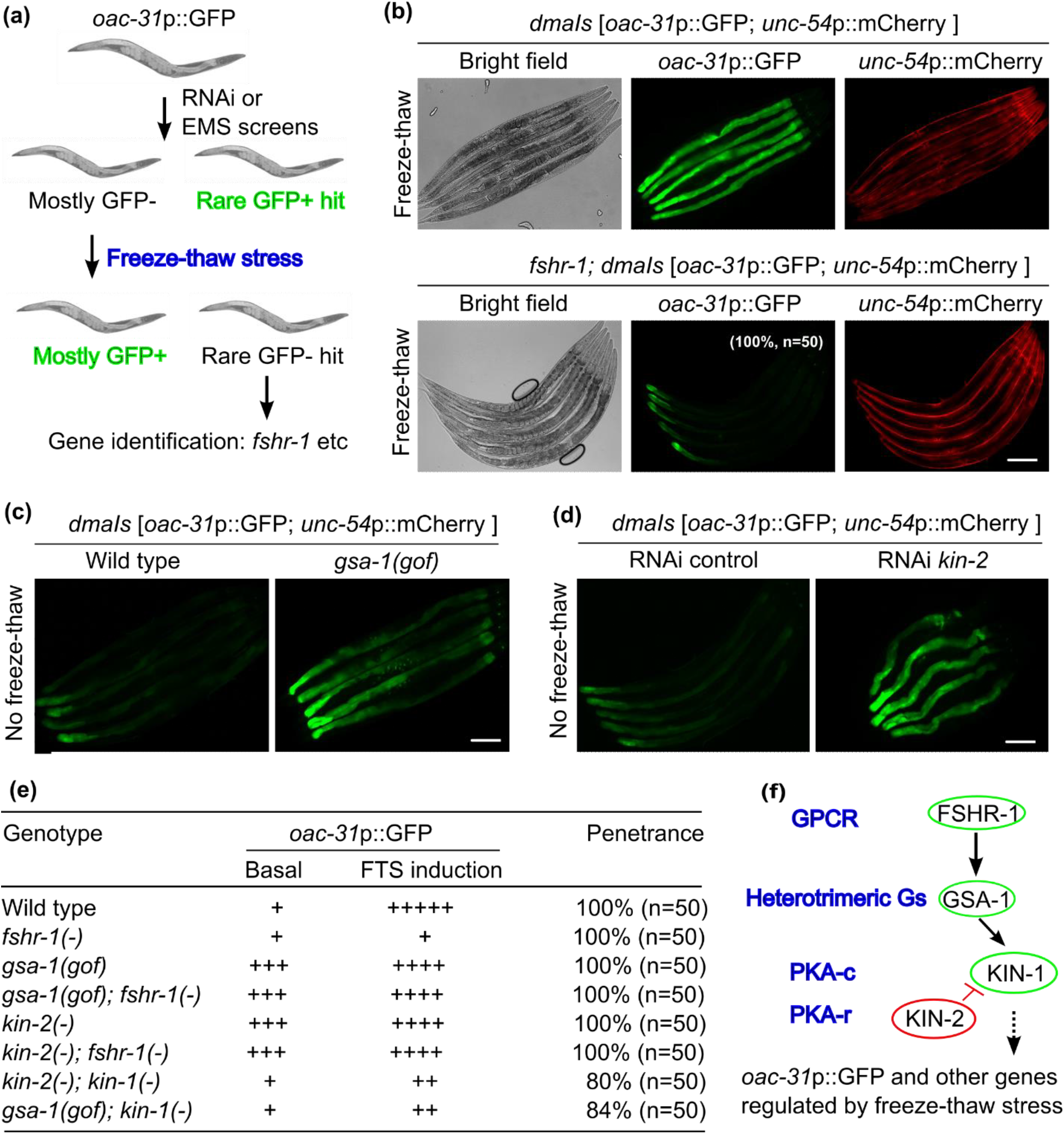
Genetic screens identify FSHR-1/GPCR signaling in regulating freeze-thaw induced *oac-31*. **(a).** Schematic of RNAi and EMS screens. **(b)**, Representative epifluorescence images showing FTS induction of *oac-31*p::GFP is blocked in *fshr-1* deletion mutants. **(c)**, Representative epifluorescence images showing constitutive induction of *oac-31*p::GFP in *gsa-1* gain-of-function mutants without FTS. **(d)**, Representative epifluorescence images showing constitutive induction of *oac-31*p::GFP in *kin-2* RNAi-treated animals without FTS. **(e)**, Summary table listing the *oac-31*p::GFP phenotype (with the numbers of “+” signs qualitatively indicating the GFP fluorescence levels) and penetrance of FTS induction in animals with genotypes indicated. **(f)**, Schematic of the FSHR-1/GSA/PKA pathway in regulating *oac-31* expression in response to FTS, based on genetic epistasis analysis. Scale bar, 100 μm.

*fshr-1* encodes the sole ortholog of glycoprotein hormone receptor family proteins in the *C. elegans* genome (Cho et al. 2007; Kudo et al. 2000). FSHR-1 family proteins are GPCRs that couple to the small G-protein Gαs to stimulate cAMP production and downstream protein kinase A (PKA) signaling. We found that a gain-of-function (GOF) mutation (Schade et al. 2005) in the *C. elegans* Gαs gene *gsa-1* strongly activated *oac-31*p::GFP in the intestine even without FTS (Figure 2c). GOF of the Gαs GSA-1 leads to cAMP elevation and activation of PKA, which comprises the catalytic subunit KIN-1 and the inhibitory regulatory subunit KIN-2 in *C. elegans* (Lu et al. 1990). We found that *kin-2* RNAi led to constitutive *oac-31*p::GFP activation as *gsa-1* GOF did (Figure 2d), and both phenotypes can be suppressed by *kin-1* but not *fshr-1* RNAi (Figure 2e). These results indicate that FTS up-regulates *oac-31* via a classic FSHR-1/GPCR-mediated cAMP and KIN-1/PKA signaling cascade (Figure 2f).

### FSHR-1 acts in the intestine to regulate ZIP-10 and cold-induced phenoptosis

We next examined the site of expression/action of FSHR-1 and its physiological consequences in FTS-induced phenoptosis. A transcriptional reporter with GFP driven by the promoter of *fshr-1* revealed its predominant intestinal site of expression (Figure 3a). A translational reporter with mCherry driven by the promoter and the genomic protein-coding sequence of *fshr-1* also revealed an intestinal pattern of FSHR-1::mCherry (Figure 3b). Transgenic wild-type *fshr-1* driven by the intestine-specific *ges-1* promoter rescued *oac-31* induction by FTS in *fshr-1* mutants, whereas RNAi against *fshr-1* specifically in the intestine nearly abolished *oac-31* induction by FTS (Figure 3c). FTS-induced *oac-31* also requires *zip-10* (Figure 1f), consistent with its role in mediating transcriptional responses to cold stress in intestinal cells. Furthermore, we found that FTS strongly induced *zip-10* expression itself in control RNAi but not intestine-specific *fshr-1* RNAi-treated animals (Figure 3d). Although FTS transcriptionally up-regulates *zip-10* through FSHR-1, high baseline expression of ZIP-10 likely also contributes to *oac-31* expression given its rapid induction kinetics. These results indicate that FSHR-1 acts in the intestine to promote FTS-induced *zip-10* and *oac-31* gene expression.

**Figure 3:**
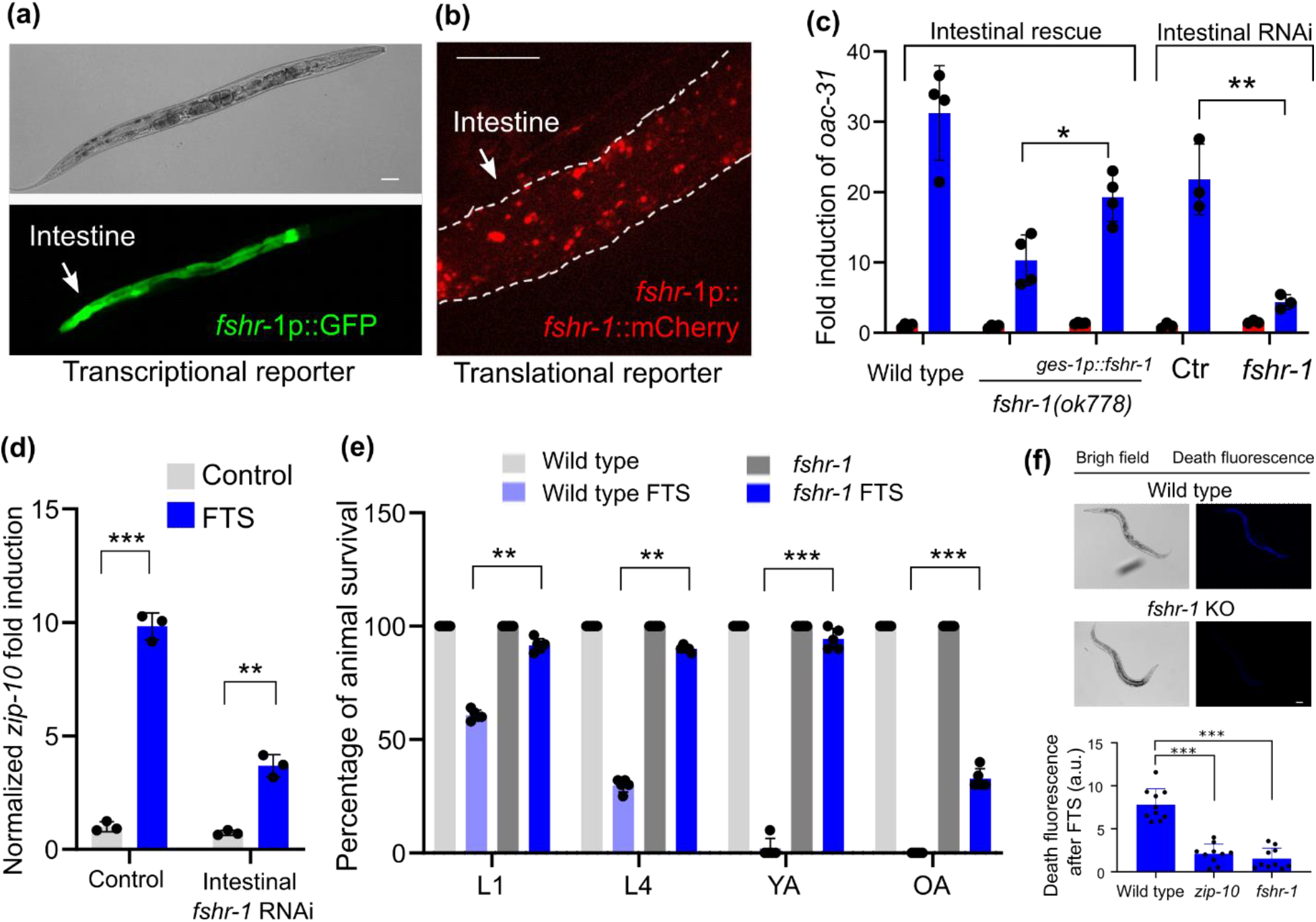
*fshr-1* is expressed and acts in the intestine to regulate FTS induction of *oac-31* and phenoptosis. **(a)**, Representative epifluorescence image of the transcriptional *fshr-1*p::GFP reporter showing predominant expression of GFP in the intestine. **(b)**, Representative epifluorescence image of the translational *fshr-1p::fshr-1*::mCherry reporter showing intestinal expression pattern. **(c)**, Tissue-specific rescue (*fshr-1* cDNA driven by the intestine-specific *ges-1* promoter) and *fshr-1* RNAi (in the strain *rde-1(ne219); Is[Pges-1::RDE-1::unc54 3’UTR; Pmyo2::RFP3]*) analysis showing *fshr-1* acting primarily in the intestine for FTS induction of *oac-31*. Values are means ± S.D with \*\**P* < 0.01 and \**P* < 0.05 (two-way ANOVA for genotype-effect interaction and post-hoc Tukey HSD, N > 3 independent experiments, n > 50 for each experiment). **(d)**, qRT-PCR results showing induction of *zip-10* by FTS was attenuated in animals with intestine-specific *fshr-1* RNAi. Values are means ± S.D with **P < 0.01, \*\*\**P* < 0.001 (two-way ANOVA for genotype-effect interaction and post-hoc Tukey HSD, N = 3 independent experiments, n > 50 for each experiment). **(e)**, Quantification of phenoptosis based on percentages of larval (L1 and L4) and adult (YA: 48 hrs post L4, OA: 5 days post L4) stage animals survived after FTS treatment. Values are means ± S.D with \*\**P* < 0.01 and \*\*\**P* < 0.001 (two-way ANOVA for genotype-effect interaction and post-hoc Tukey HSD, N > 3 independent experiments, n > 50 for each experiment). **(f)**, Representative bright field and fluorescence images showing characteristic blue “death fluorescence” in wild type but not *zip-10* or *fshr-1* mutants (\*\*\**P* < 0.001, one-way ANOVA with post-hoc Tukey HSD, N = 10) 24 hrs after FTS. Scale bar, 100 μm.

We next assessed the role of the FSHR-1 signaling cascade in FTS-induced phenoptosis. Compared to wild type, *fshr-1* loss-of-function mutants showed markedly increased rates of survival after FTS for both larvae and adults (Figure 3e). We observed death-characteristic blue fluorescence after the FTS treatment in wild-type adult animals as previously reported for naturally dying animals (Coburn et al. 2013), but markedly less so in *zip-10* or *fshr-1* mutants (Figure 3f). In addition, FLR-2 is the sole *C. elegans* ortholog of glycoprotein hormones and putative FSHR-1 ligand (Oishi et al. 2009, p.2) that may signal through the FSHR/GSA-1/PKA cascade (Figure 2). Like *fshr-1* deficient animals, loss of *flr-2* or *kin-1* caused defective *oac-31* induction by FTS (Figure 4a). To determine the epistatic relationship of the genes in the FLR-2/FSHR-1 pathway for FTS-induced phenoptosis, we examined how their loss-of-function or gain-of-function impacted FTS-induced phenoptosis at L4 stages, in which either positive or negative effects on phenoptosis can be quantitatively measured (Figure 4b). Both *flr-2* single and *flr-2; fshr-1* double loss-of-functions (by mutant or RNAi) showed defective

**Figure 4:**
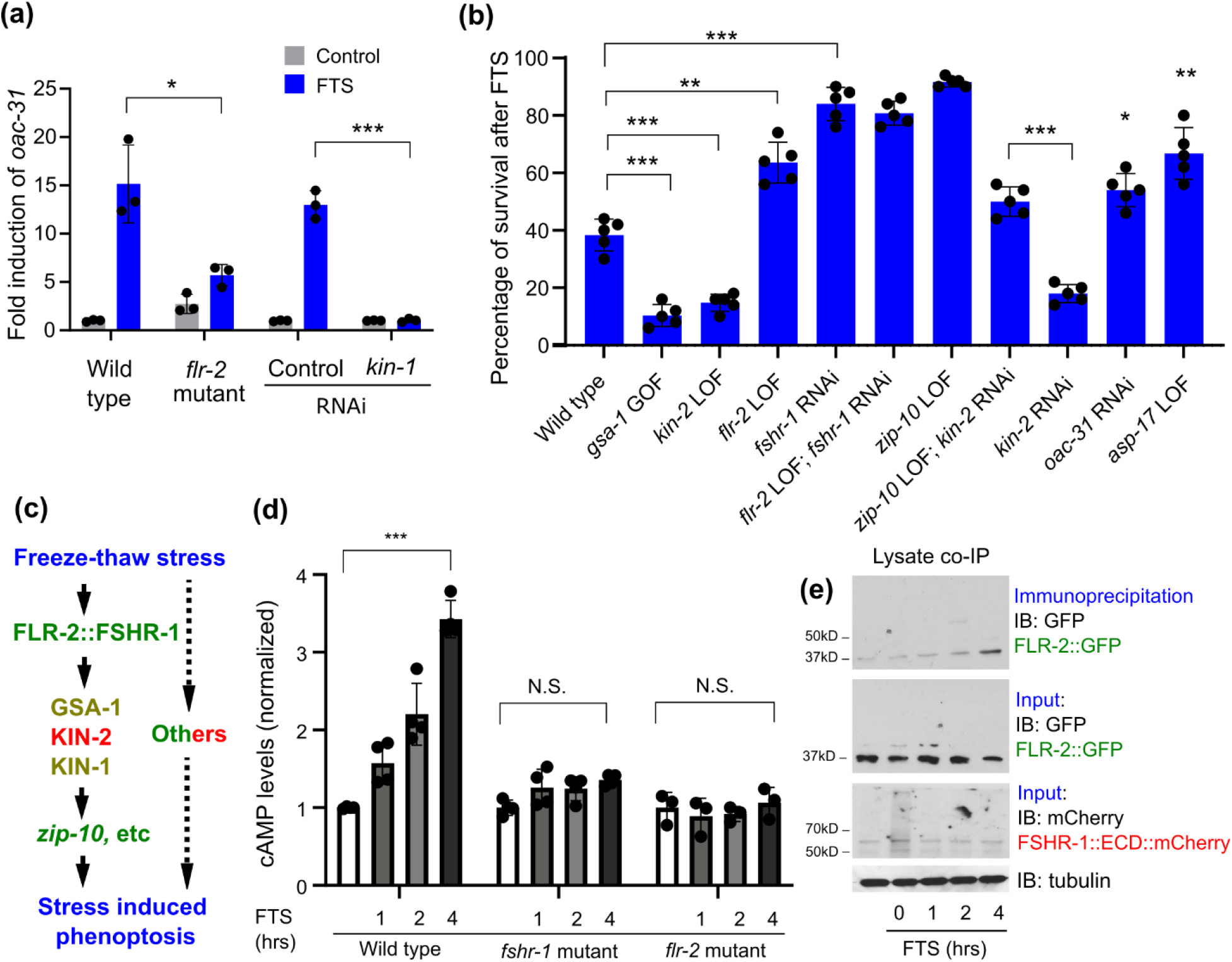
FLR-2/FSHR-1-mediated GPCR/cAMP signaling drives phenoptosis. **(a)**, qRT-PCR results showing abolished *oac-31* induction by FTS in *flr-2* reduction-of-function mutants, mimicking *fshr-1* mutants. \*\**P* < 0.01 (two-way ANOVA for genotype-effect interaction and post-hoc Tukey HSD, N = 3 independent experiments, n > 50 for each experiment). **(b)**, Table summary of percentages of larval L4-stage animals with indicated genotypes survived after FTS treatment \**P* < 0.05 and \*\*\**P* < 0.001 (one-way ANOVA and post-hoc Tukey HSD N = 5 independent experiments, n > 50 for each experiment). **(c)**, Schematic of the genetic pathway leading to FTS-induced phenoptosis based on epistasis and gene expression analysis. Known genes or proteins promoting (green) or antagonizing (red) cold stress-induced phenoptosis are noted and with other contributors (dashed line). **(d)**, Normalized ELISA results showing FTS-induced (no FTS control, 1 hr, 2hrs, 4 hrs of recovery after freezing) elevation of cAMP was abolished in *flr-2* and *fshr-1* mutants. Values are means ± S.D with \*\*\**P* < 0.001 (two-way ANOVA for genotype-effect interaction and post-hoc Tukey HSD, N > 3 independent experiments, n > 50 for each experiment). N.S., non-significant. **(e)**, Representative western blots of GFP-tagged FLR-2, mCherry-tagged FSHR-1::ECD(extracellular domain)::mCherry from input and co-immunoprecipitation lysate samples showing time-dependent (no FTS control, 0 hr, 1 hr, 2 hrs, and 4 hrs of recovery after freezing) binding of FLR-2::GFP and FSHR-1::ECD::mCherry upon FTS treatment (tubulin as loading control).

FTS-induced phenoptosis (Figure 4b). Consistent with their roles in regulation of *oac-31* downstream of FSHR-1, *gsa-1* gain-of-function or *kin-2* loss-of-function exhibited enhanced FTS-induced phenoptosis, while such enhancement can be suppressed with *zip-10* loss-of-function mutations (Figure 4b). Loss of ZIP-10 strongly attenuated FTS-induced phenoptosis (Figure 1a), while individual loss of its target genes *oac-31* or *asp-17* (previously identified as a ZIP-10-dependent gene encoding a protease) decreased FTS-induced phenoptosis to a lesser extent (Figure 4b). These results indicate that FTS promotes phenoptosis through an FSHR-1/GSA-1/PKA/ZIP-10 cascade (Figure 4c).

FSHR-1 has been previously implicated in responses to oxidative and pathogen stresses (Miller et al. 2015, p.1; Powell et al. 2009, p.). Among the membrane lipids that can be oxidized, cholesterol is most prone to oxidation by reactive oxygen species (Murphy & Johnson 2008). Consistent with previous studies supporting a role of FSHR-1 in protecting against oxidative stress associated with cell membrane lipids, we found that *fshr-1* RNAi caused higher sensitivity than wild type to exposure of exogenous cholesterol when supplemented with the oxidizing agent Paraquat (Figure S3a). In addition, we found that the cold stress or FTS resilience phenotypes caused by RNAi against *fshr-1* can be modulated by the amount of exogenous (supplemented in media) or endogenous (manipulated by *ncr* expression) cholesterol availability (Figure S3b, c, d). These results suggest that resilience to severe hypothermic stress can come with a physiological trade-off in organismic resilience to cholesterol oxidative stresses.

We next explored how FTS may regulate FLR-2/FSHR-1. Based on RNAseq results, FTS did not affect expression levels of *flr-2* or *fshr-1* (Figure S4a). While *flr-2* is expressed predominantly in neurons based on single-cell RNA results (Taylor et al. 2021), we generated a translational reporter for FLR-2 fused with GFP and identified its localization to coelomocytes, which likely uptake secreted forms of FLR-2::GFP from neurons (Figure S4b). However, *flr-2* reporters showed no apparent changes after FTS treatment (Figure S4b). These data suggest that neuron-derived systemically-acting FLR-2 is required for FTS induction of *oac-31* and phenoptosis, but not directly regulated by FTS transcriptionally. Nonetheless, we observed that FTS induced marked elevation of intracellular cAMP levels in wild type but not *flr-2* or *fshr-1* mutants (Figure 4d). In addition, co-immunoprecipitation experiments using transgenic animals show that GFP-tagged FLR-2 can increasingly bind to mCherry-tagged FSHR-1 extracellular domains after FTS in a time-dependent manner (Figure 4e), consistent with an increase in cAMP levels after FTS. As glycoprotein hormones typically bind to its obligatory GPCRs to stimulate cAMP levels, these results further support the notion that FSHR-1-dependent GPCR signaling actively mediates response to FTS, rather than plays a permissive role, to drive stress-induced gene expression and phenoptosis.

### A mathematical model to explain plausible adaptive functions of phenoptosis

At first glance, existence of phenoptosis and a genetic pathway promoting phenoptosis would seem incompatible with the evolutionary theory as genes promoting phenoptosis would compromise individual fitness, thus not be subject to selection pressure. Why would a genetic program possibly function to promote phenoptosis? We established a mathematical model (Figure 5) built upon the evolutionary concept of kin selection that may explain adaptive or beneficial functions of phenoptosis at the population level. We define three key parameters in the model that determine the growth of reproduction-competent haploid genomes in a population, including the parameter of α (aging rate), β (genome replication rate), and μ (cohort competition/infection cost after stress). More stress sensitivity (i.e. phenoptosis susceptible) in weak or old (v.s. fit or young) animals would decrease μ, since weak or old animals contribute less than fit or young animals in the growth of replication-competent genomes in the population. Simulated results from our model assuming constant parameters in a given population show that the number of healthy individuals in a starting population can increase faster than a population without stress-inducible phenoptosis, under conditions where there is competition for limited resources to reproduce or infectious spreading of pathogens among individuals (Figure 5). According to the model, stress-induced phenoptosis may act against the unfit group members (severely stressed adults less reproductive or sick individuals shedding infectious pathogens) to benefit the whole group at the population level.

**Figure 5:**
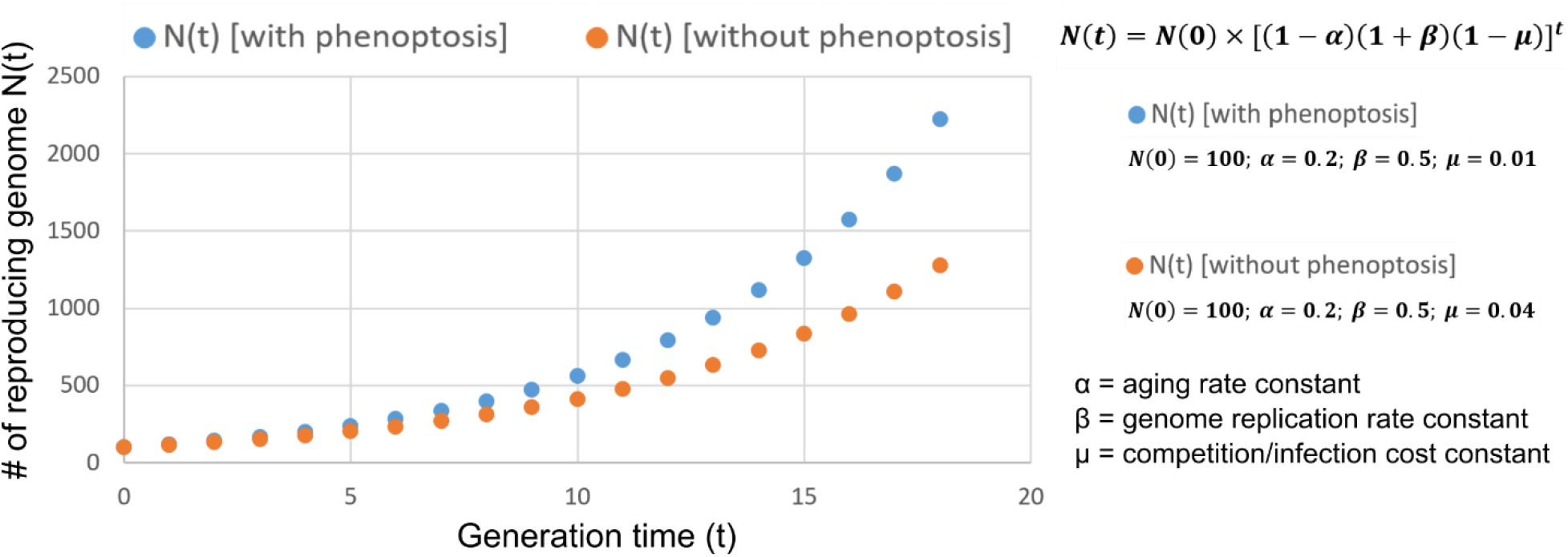
A simplified mathematical model to simulate potential adaptive function of phenoptosis. Shown is a graph simulating proliferation of reproducing genomes following the equation ***N*(*t*) = *N*(0) × [(1 – *α*)(1 + *β*)(1 – *μ*)]^t^**, where N (0) = Initial number of reproduction-competent haploid genomes in a population, N (t) = Number of reproducing haploid genomes in a population at the generation time of t, α = aging rate constant, β = genome replication rate constant, μ = cohort competition/infection cost constant. For simulation, we used ***N*(0) = 100; *α* = 0.2; *β* = 0.5; *μ* = 0.04** for the growth of numbers of reproducing genomes without phenoptosis or decreased phenoptosis (e.g. in *fshr-1* mutants), while ***N*(0) = 100; *α* = 0.2; *β* = 0.5; *μ* = 0.01** with phenoptosis (e.g. in the wild-type *C. elegans*). For simplicity, we make assumptions on the clonal population structure and growth characteristics in a resource-limiting competitive environment and that α and β remain constant. Note that μ is lower with increased phenoptosis given limited resources for a population, since increased stress-induced phenoptosis in less reproduction-competent individuals leads to reduced competition for resources or vulnerability to pathogen infection at the population level.

## DISCUSSION

While genes can control how cells die in well-known processes including apoptosis, necroptosis, and pyroptosis (Shi et al. 2017; Metzstein et al. 1998; Weinlich et al. 2017), how genes might control the death of an organism remains largely unclear. Based on comprehensive genetic analyses and stress-responding organismic phenotypes, we propose a model in which FSHR-1 signaling may mediate FTS-induced phenoptosis in *C. elegans*. After exposure to severe hypothermic stress conditions, FLR-2 may promote FSHR-1 activation and signaling through a GPCR-dependent Gαs-cAMP-PKA pathway, leading to increased expression of *zip-10, oac-31* and other genes in a genetic program that coordinately drive phenoptosis. Site-of-expression and site-of-action analyses indicate that this GPCR pathway operates in the intestine, consistent with intestinal expression and cell-autonomous regulation of *zip-10* and *oac-31*. Regarding the mechanism of how FTS activates FLR-2/FSHR-1, it is conceivable that sensory neurons may detect thermal stress signals to promote release of FLR-2 to act systemically, while intestinal cells may also adjust membrane properties upon FTS to modulate FLR-2/FSHR-1 activation (Ernst et al. 2016). Although ZIP-10 is essential for up-regulation of many genes induced by FTS, mechanisms linking FSHR-1/GPCR, KIN-1/PKA to *zip-10* regulation remain unidentified. Candidate factors may include CREB/bZIP family transcription factors (Lakhina et al. 2015; Zhang et al. 2017, p.1) and the p38/JNK pathways (Zhang et al. 2017; Hattori et al. 2013). In particular, the stress-responding *kgb-1/JNK* pathway has been shown to promote detrimental effects during *C. elegans* aging (Twumasi-Boateng et al. 2012). Whether and how KGB-1 might play similar roles in FTS-induced phenoptosis remain to be investigated in future studies.

Various types of environmental stress other than FTS have been shown to trigger acute organismic death in *C. elegans* that is preceded by cellular necrosis in the intestine (Ezcurra et al. 2018; Coburn et al. 2013; Zhang et al. 2016; Luke et al. 2007). Mediators linking stress signals to cellular necrosis programs remain largely undefined and are likely stress type-specific. As ZIP-10 appears both required and partially sufficient to drive cold-induced phenoptosis, it may represent a master transcriptional regulator downstream of stress-sensing GPCR signaling, leading to systemic necrosis and organismic death through its target gene effectors, including those encoding proteases we previously identified (Jiang et al. 2018). Our results suggest that the *ISY-1/mir-60* axis plays a gatekeeping role for ZIP-10 activation and phenoptosis, whereas the FLR-2/FSHR-1/GSA/KIN-2 axis is likely instructive by linking FTS to genes regulating cellular necrosis or promoting organismic sensitivity to lethal effects of FTS. It is worth noting that the death-promoting effect of PKA signaling after FTS differs from its protective effect under constitutive cold conditions (Liu et al. 2017), underscored by the fact that strong induction of the *zip-10* transcriptional program requires the warming or recovery phase post cold or freezing stresses (Jiang et al. 2018). FSHR-1 and ZIP-10 have also been shown to mediate host cell responses to oxidative stress, various types of pathogens and immune aging (Powell et al. 2009; Miller et al. 2015, p.1; Lee et al. 2021, p.1; Kato et al. 2016). Indeed, we found that *fshr-1* loss-of-function mutants, while being more FTS resistant than wild type, were less tolerant of cholesterol oxidative stress (Figure S4). How FSHR-1/GPCR, KIN-1/PKA or ZIP-10/bZIP may function in other stress paradigms similarly in promoting organismic phenoptosis or play context-specific roles in physiological trade-off awaits further studies.

We note that *oac-31*, like several other *oac* genes (*oac-3, 6, 14, 20*) strongly up-regulated by FTS (Figure S1g), encodes a *C. elegans* homolog of sterol O-acyltransferase (SOAT). SOAT converts accessible cholesterol into ester forms for storage in lipid droplets (Luo et al. 2020; Sevanian & Peterson 1986). While the biochemical function of OAC-31 remains further characterized, a recent study showed that cold shock can promote lipid movement from the intestine to the germline (Gulyas & Powell 2021). Interestingly, natural aging of *C. elegans* is accompanied with autophagy-mediated conversion of intestinal biomass into yolk, contributing to late-life aging pathologies and mortality (Ezcurra et al. 2018). Thus, it is plausible that FSHR-1-dependent up-regulation of SOAT may be part of a genetic program that converts somatic biomass into nutrient/energy reserves for trafficking to the germline to benefit the progeny at the expense of parental death. In early life, such genetic programs may have evolved to promote reproductive health, while exhibiting antagonistic pleiotropic effects promoting organismic death and aging in late life. Phenoptosis-promoting effects of FSHR-1/ZIP-10 are also dependent on developmental stages, stress types and severity, reflecting on their pleiotropic context-dependent nature. At the population level, the phenoptosis of severely stressed weak or sick adults (aged or infected) may also be beneficial as limiting resources can be funneled to less severely stressed and reproductively active younger individuals. These two scenarios (“disposable soma” and “kin selection”) are not mutually exclusive and may have been subject to selection pressure and explain the conservation of phenoptosis and its pathway of regulation in certain species, particularly those with clonal population structures (Galimov et al. 2019). Together, our findings identify key signaling mediators of severe cold stress-induced phenoptosis in *C. elegans* and provide empirical evidence of how phenoptosis can be genetically regulated with plausible evolutionary significance.

## MATERIALS AND METHODS

### *C. elegans* strains

*C. elegans* strains were maintained with standard procedures unless otherwise specified. The N2 Bristol strain was used as the reference wild type, and the polymorphic Hawaiian strain CB4856 was used for genetic linkage mapping and SNP analysis (Brenner 1974; Davis et al. 2005). Forward genetic screen for *oac-31*p::GFP mutants after ethyl methanesulfonate (EMS)-induced random mutagenesis was performed as described previously (Ma et al. 2015; Ma et al. 2012). Feeding RNAi was performed as previously described (Kamath & Ahringer 2003). Customized sub libraries of RNAi clones were constructed by manual selection from the main Ahringer library based on gene expression abundance (TPM>2) in the intestine and encoded proteins as putative transmembrane receptors, signaling proteins and transcription factors (Table S2). Transgenic strains were generated by germline transformation as described (Mello et al. 1991). Transgenic constructs were co-injected (at 10 - 50 ng/μl) with dominant *unc-54p::mCherry* or *rol-6* markers, and stable extrachromosomal lines of mCherry+ or roller animals were established. Genotypes of strains used are: *gsa-1(ce81) I, ncr-2(nr2023) III, fshr-1(ok778) V, flr-2(ut5) V, ncr-1(nr2022) X. dmaEx616 [fshr-1p::GFP; unc-54p::mCherry], vjEx1449[ges-1p::fshr-1], dmaEx617 [fshr-1p::fshr-1::GFP; unc-54p::mCherry], dmaEx620 [rpl-28p::fshr-1ECD::mCherry], rde-1(ne219); Is[Pges-1::RDE-1::unc54 3’UTR; Pmyo2::RFP3]* (gift of Dr. Meng Wang’s laboratory)*, dmaIs136 [flr-2p::flr-2::GFP], dmaIs117 IV [oac-31p::GFP; unc-54p::mCherry]*.

### Sample preparation for RNA sequencing and data analysis

Control N2 animal were maintained at 20 °C. For freeze-thaw stress, N2 animals were exposed to −20 °C for 25 mins followed by 1 hr recovery at 20 °C. Upon sample collection, the animals were washed down from NGM plates using M9 solution and subjected to RNA extraction using the RNeasy Mini Kit from Qiagen. 1 μg total RNA from each sample was used for sequencing library construction. RNA-seq Library preparation and data analysis were performed as previously described (Jiang et al. 2018). Three biological replicates were included for each treatment. The libraries were sequenced to 51 bp at single end by the Center for Advanced Technology (CAT) of the University of California, San Francisco using a Hiseq-3000 system. The cleaned RNAseq reads were mapped to the genome sequence of *C. elegans* using hisat2 (Kim et al. 2015). The mapped reads were assigned to the genes using featureCounts (Liao et al. 2014). Abundance of genes was expressed as RPKM (Reads per kilobase per million mapped reads). Identification of differentially expressed genes was performed using the DESeq2 package (Love et al. 2014).

### Quantitative RT-PCR

25 μl pellet animals were resuspended in 400 μl lysis buffer of Quick-RNA MiniPrep kit (Zymo Research, R1055) then lysed by TissueRuptor (Motor unit ‘8’ for 1 min). Total RNA was extracted following the instruction (Zymo Research, R1055). 1 μg RNA/sample was reverse transcribed into cDNA (Thermo Fisher Scientific, 18080051). Real-time PCR was performed by using Roche LightCycler96 (Roche, 05815916001) system and SYBR Green (Thermo Fisher Scientific, FERK1081) as a dsDNA-specific binding dye. qRT-PCR condition was set to 95°C for denaturation, followed by 45 cycles of 10 s at 95°C, 10 s at 60°C, and 20 s at 72°C. Melting curve analysis was performed after the final cycle to examine the specificity of primers in each reaction. Relative mRNA was calculated by ΔΔCT method and normalized to actin. Primers for qRT-PCR: *oac-31* (Forward, ATGGCTTAAATCGCTGGAAA; Reverse, ATCTTTCGCCATCAATACGG), *W09G12.7*, Forward, CGAAGATCACTCGAACACGA; Reverse, CCATTTTTACGAAGGCTGGA), *tsp-1* (Forward, CTTTGATTGCCGTTGGATTT; Reverse, CCCAAAGAAAGGCCGATAAT), *act-3* (Forward, TCCATCATGAAGTGCGACAT; Reverse, TAGATCCTCCGATCCAGACG).

### Freezing resilience assay

Animals were cultured under non-starved conditions for at least 4 generations at 20 °C before cold and freezing resilience assay. For freezing resilience assay, bleach-synchronized L1, L4, young (48 hrs post L4) and older adult (5 days post L4) populations were kept at −20 °C for 45 mins and then recovered for 24 hrs at 20 °C. For such experiments, NGM plates spread with equal agar thickness seeded with equal amounts of OP50 were used while hypothermic temperature readings were monitored by thermometers to ensure minimal fluctuation. After freezing shock, animals were moved to 20 °C for recovery and scored as dead if they showed no pumping and movement upon light touch with the body necrosis subsequently confirmed.

### cAMP assay

N2, *fshr-1* and *flr-2* mutants were freeze shocked (−20 °C) for 25 mins, followed by recovery at 20 °C for 0, 1, 2 or 4 hrs. Animals were harvested and washed three times with M9 and 20 μl pellet animals were lysed directly in ELISA lysis buffer. Then worm extracts were collected by centrifugation. The cAMP levels in the supernatant were determined using the cyclic AMP ELISA kit (Cayman Chemical, Ann Arbor, MI) according to the manufacturer’s instructions. BCA kit was used to measure the concentration of protein as normalizing control.

### Co-immunoprecipitation assay

Transgenic animals expressing *flr-2p::flr-2::GFP* and *fshr-1::ECD::mcherry* were cultured under non-starved conditions at 20 °C on NGM plates before freezing treatment. For freezing treatment, mixed stage worms were kept at −20 °C for 25 mins and then recovery for 0, 1, 2, 4 hrs at 20 °C. Upon sample collection, the animals were washed down from NGM plates using M9 solution and subjected to protein extraction using ice-cold 2x lysis buffer (60 mM HEPES, pH = 7.4, 100 mM potassium chloride, 0.1% Triton X, 4 mM magnesium chloride, 10% glycerol, 2 mM DTT with RNase inhibitor, protease inhibitor, and phosphatase inhibitors) and lysed with sonication. The lysate was then centrifuged at 15,000 rpm for 15 mins to remove cell debris. The supernatant was incubated with 4 μl of pre-clean magnetic beads at 4 °C for 15 minutes. To precipitate *fshr-1*::ECD::mcherry proteins, the sample was placed into a 1.5 ml RNase-free tube containing RFP-trap magnetic beads at 4 °C overnight. The beads were then washed with 400 μl of lysis buffer for 5 times and incubated with *flr-2::flr-2::GFP* lysate samples at 4 °C overnight. The beads were then washed with 400 μl of lysis buffer for 5 times and resuspended in 40 μl of 1 ×Laemmli Sample Buffer. The samples were then boiled at 95 °C for 10 minutes and subject to Western blot analysis.

### Statistical analysis

Data were analyzed using GraphPad Prism 9.2.0 Software (Graphpad, San Diego, CA) and presented as means ± S.D. unless otherwise specified, with *P* values calculated by unpaired two-tailed t-tests (comparisons between two groups), one-way or two-way ANOVA (comparisons across more than two groups) followed by Tukey’s post-hoc tests and adjusted with Bonferroni’s corrections.

## Data availability

The RNAseq read datasets were deposited in NCBI SRA (Sequence Read Archive) under the BioProject accession PRJNA763790. All other data newly generated for this study are included in this article.

## Acknowledgments

Some strains were provided by the CGC, which is funded by NIH Office of Research Infrastructure Programs (P40 OD010440), and from Drs. Derek Sieburth and Meng Wang laboratories. The work was supported by NIH grant 1R35GM139618 and the Packard Fellowship in Science and Engineering (D.K.M) and National Natural Science Foundation of China, Grant No. 32271165, 31971077 and 31971047 (C.Z., C.W.).

## Author contributions

C.W., B.W., C.Z. and D.K.M. designed, performed and analyzed the experiments and wrote the manuscript. Y.L. performed RNA sequencing and analysis. C.Z. and D.K.M. provided resources and supervised the project.

## Competing interests

The authors declare no competing interests.

## Materials & Correspondence.

Correspondence and material requests should be addressed to Dengke K. Ma, Ph.D.

---

**Figure S1:**
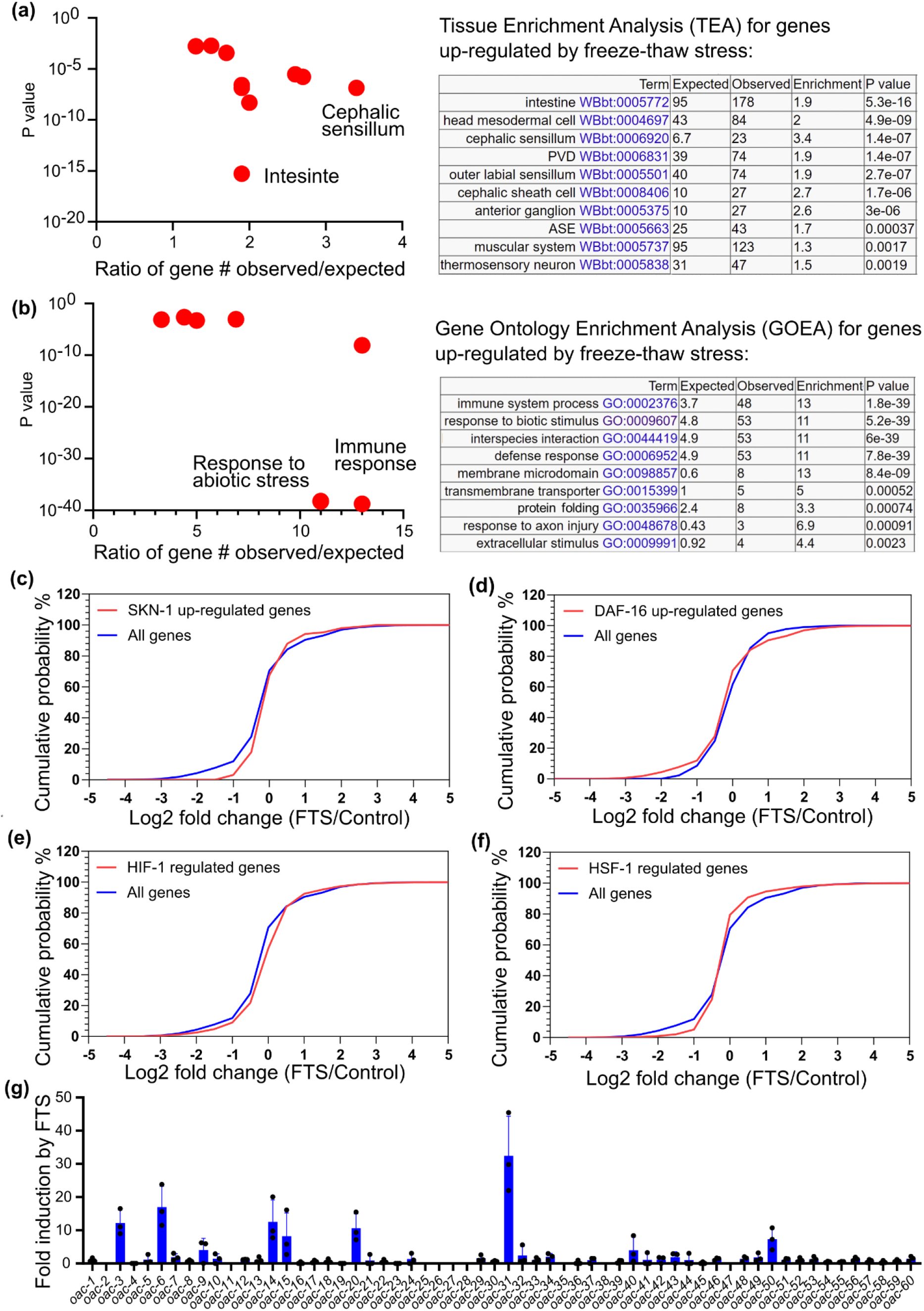
Transcriptome profiling and analysis of genes up-regulated by FTS. **(a)**, Tissue Enrichment Analysis (TEA) of genes up-regulated by FTS showing intestine as a major site of gene regulation. **(b)**, Gene Ontology Enrichment (GOE) Analysis of genes up-regulated by FTS showing abiotic stress and immune response as major processes involved. **(c-f)**, Cumulative probability analysis of genes regulated by SKN-1 (**c**), DAF-16 (**d**), HIF-1 (**e**), HSF-1 (**f**) (Brunquell et al. 2016, p.1; Kumar et al. 2015; Oliveira et al. 2009; Steinbaugh et al. n.d.) does not show marked enrichment of those also regulated by FTS (NCBI Sequence Read Archive BioProject accession PRJNA763790). **(g)**, Normalized fold induction of gene expression for the entire *oac* gene family based on RNAseq results, showing *oac-31* as the most up-regulated family member.

**Figure S2:**
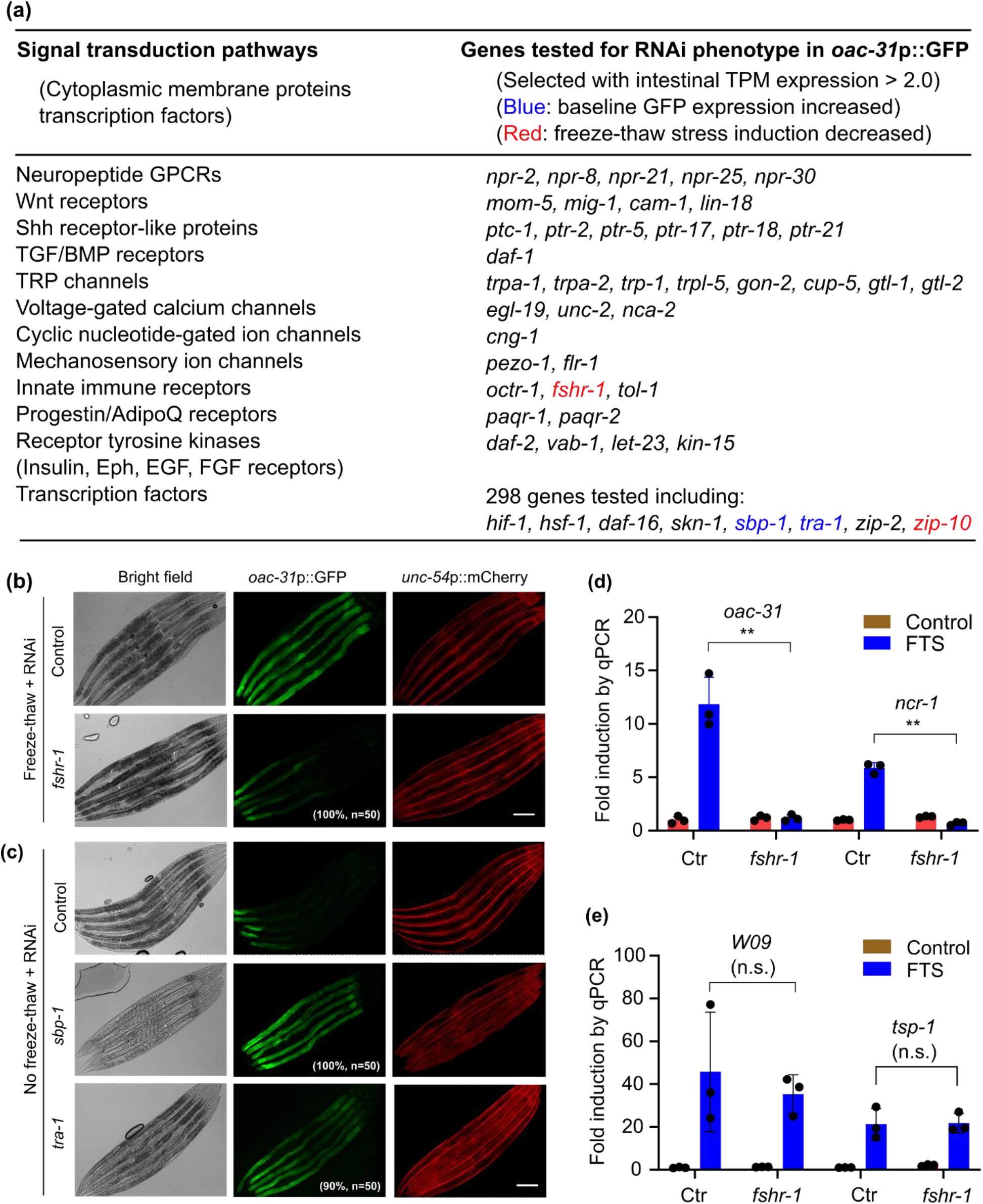
RNAi screens identify genes that regulate *oac-31*p::GFP induction by FTS. **(a)**, Table listing RNAi clones screened for *oac-31*p::GFP induction phenotype. RNAi cloned were selected based on intestinal expression and genes encoding transmembrane receptors and representing major signal transduction pathways. **(b)**, Representative fluorescence images showing suppression of FTS-induced *oac-31*p::GFP by RNAi against *fshr-1*. **(c)**, Representative fluorescence images showing constitutive activation of *oac-31*p::GFP by RNAi against *tra-1* or *sbp-1* in the absence of FTS treatment. Scale bar, 100 μm. **(d),** qRT-PCR results showing abolished induction of *ncr-1, oac-31* by FTS in animals deficient in *fshr-1*. **(e),** qRT-PCR results showing normal induction of the genes *W09G12.7* and *tsp-1*, by FTS in *fshr-1* mutants. Values are means ± S.D with \*\**P* < 0.01 and \**P* < 0.05 (two-way ANOVA for genotype-effect interaction and post-hoc Tukey HSD, N = 3 independent experiments, n > 50 for each experiment). n.s., non-significant. Scale bar, 100 μm.

**Figure S3:**
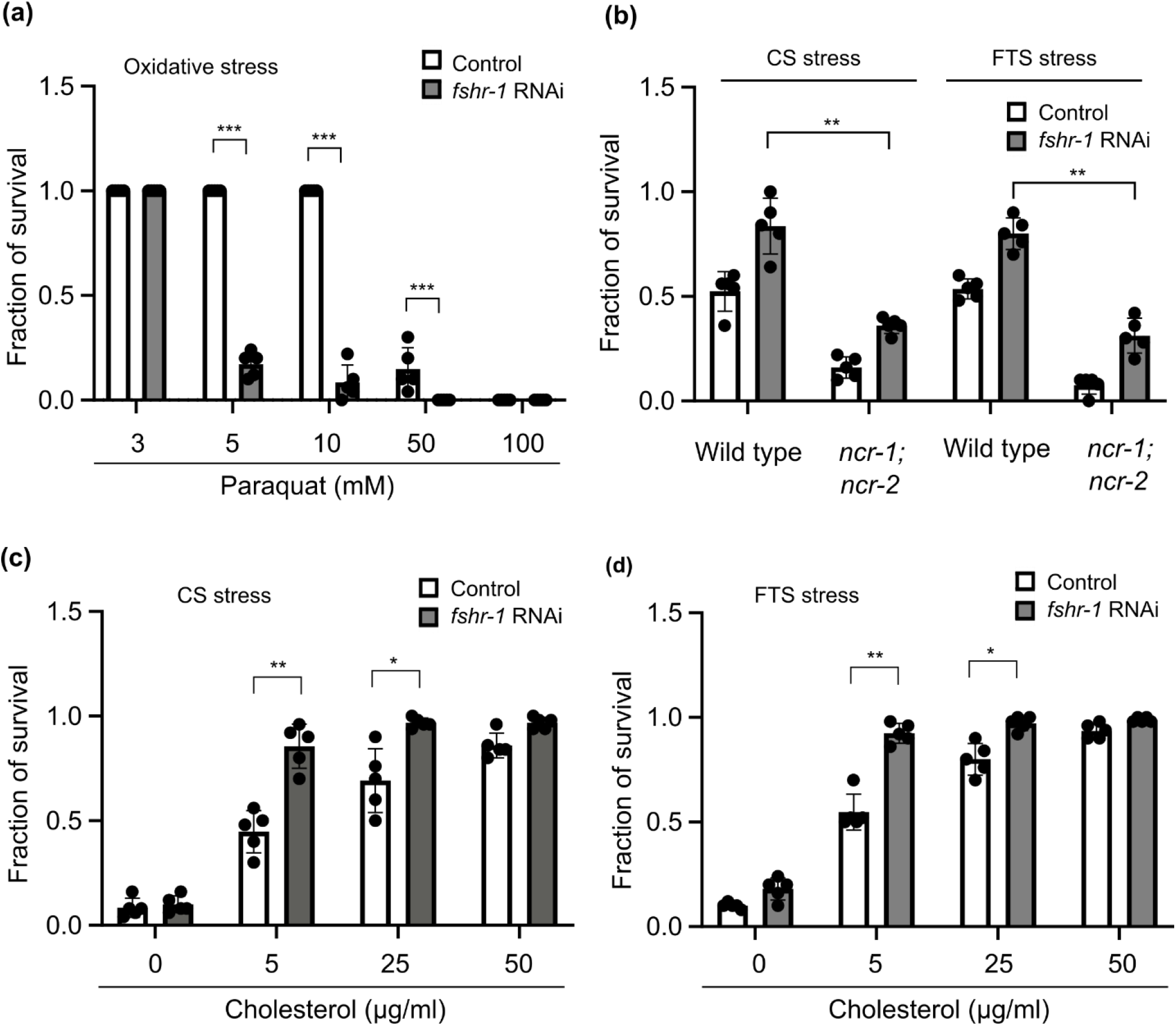
FSHR-1 regulates trade-off of organismic resilience to cholesterol oxidative and hypothermic stresses. **(a),** Survival rates of control and *fshr-1* RNAi-treated animals grown on increasing doses of Paraquat (3, 5, 10, 50 and 100 mM in NGM). **(b),** Survival rates of wild type or *ncr-1; ncr-2* double mutant animals treated with control or *fshr-1* RNAi after severe FTS (−20 °C for 45 minutes, followed by recovery for 24 hrs at 25 °C) or cold shock (CS, 4 °C for 24 hrs, followed by recovery for 24 hrs at 25 °C) grown on cholesterol (5 μg/ml). **(c),** Survival rates of control and *fshr-1* RNAi-treated animals after severe CS grown on increasing doses of cholesterol (0, 5, 25, 50 μg/ml). **(d),** Survival rates of control and *fshr-1* RNAi-treated animals after severe FTS grown on increasing doses of cholesterol (0, 5, 25, 50 μg/ml). Values are means ± S.D with \*\*\**P* < 0.001, \*\**P* < 0.01 and \**P* < 0.05 (two-way ANOVA for genotype-effect interaction and post-hoc Tukey HSD, N = 3 independent experiments, n > 50 for each experiment).

**Figure S4:**
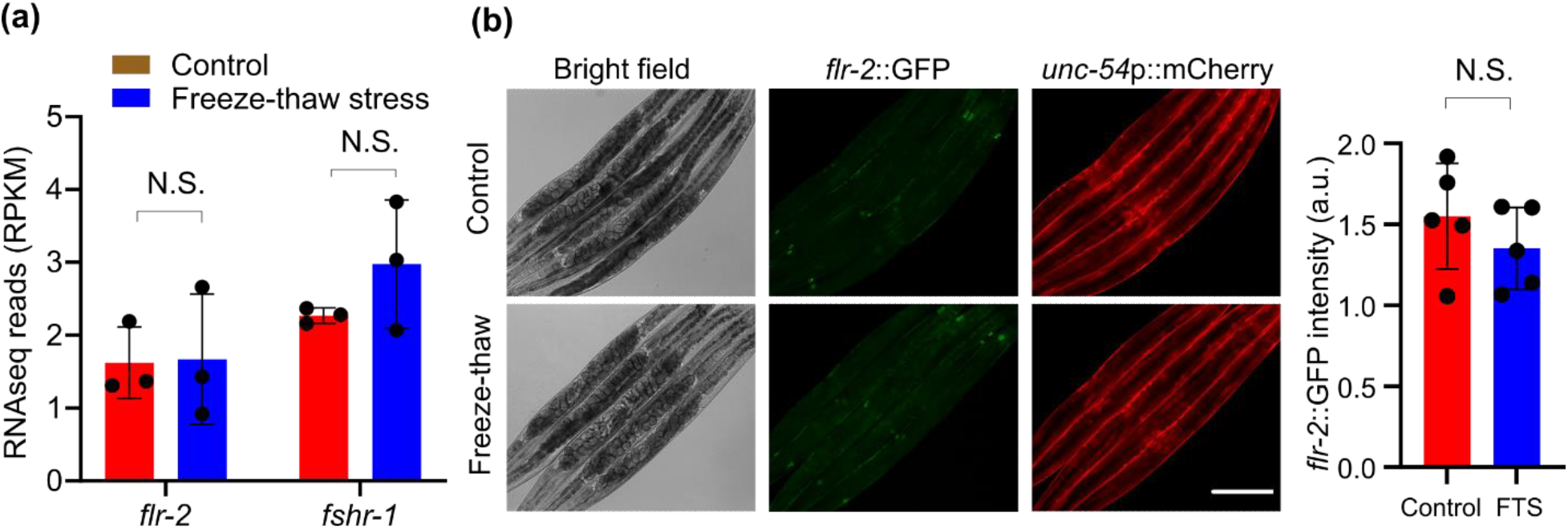
*flr-2*, which encodes the putative ligand of FSHR-1, is essential for FTS induction of *oac-31* but not transcriptionally regulated by FTS. **(a)**, RNAseq results showing no apparent up-regulation of *flr-2* or *fshr-1* by FTS (two-way ANOVA for genotype-effect interaction and post-hoc Tukey HSD, N = 3 independent experiments, n > 50 for each experiment). N.S., non-significant. **(b)**, Representative fluorescence images for the translational reporter *flr-2p::flr-2::GFP* showing its normal secretion and enrichment in coelomocytes but without apparent regulation by FTS (unpaired student-t test, N.S., non-significant).

## Notes

### Competing Interest Statement

The authors have declared no competing interest.

### Summary of Updates

Title is changed to more accurately describe findings. Discussion and introduction are expanded to richer context. More references and author info are added to reflect credit and contributions.

